# Multiparametric Correlative Topographical and Volumetric Fluorescence Microscopy

**DOI:** 10.1101/2025.07.01.662543

**Authors:** Wenzhi Hong, Ziwei Zhang, Ao Li, Ting Sun, Yunzhao Wu, Devkee M. Vadukul, Dylan Jones, Bing Li, Fengjie Liu, Francesco A. Aprile, Julia Gorelik, David Klenerman, Andrew Shevchuk

**Author notes:** **These authors contributed equally to this work.** (Wenzhi Hong, Ziwei Zhang).

## Abstract

Live-cell imaging of cell surface topography and intracellular architecture is essential for understanding cellular function. However, conventional approaches often involve trade-offs between resolution, invasiveness, and volumetric coverage. Here, we present an integrated Scanning Ion Conductance Microscope and single-objective Oblique Plane Microscope (SICM-OPM) system that enables simultaneous non-contact topographical imaging and volumetric fluorescence imaging within the same live cell. Beyond correlative live imaging, the platform supports nanomechanical mapping with tens-of-nanometres resolution, fluorescence-guided localised molecular delivery via the SICM, and benefits from reduced photobleaching due to light-sheet excitation. We demonstrate this platform’s capabilities by visualising imipramine-induced T-tubule remodelling in live cardiomyocytes, revealing subsurface detubulation while surface morphology remains preserved. Additionally, we show precision delivery of fluorescent cargos—including dextrans and α-synuclein—into diatom and mammalian cells, alongside localised stiffness mapping to evaluate mechanical responses. We believe this technique opens new avenues for correlative structural, functional, and biophysical studies in live cells, with broad relevance to cell biology, neurodegeneration, and mechanobiology.

## 1. Introduction

Direct high-resolution visualisation of cell morphology and membrane dynamics is crucial for investigating key cellular processes, such as endocytosis, lipid raft formation, receptor clustering, transmembrane transport, membrane protein trafficking, and the passage of single nanoparticles or viruses—offering essential insights into cellular function in both health and disease. However, conventional imaging techniques face significant limitations: fluorescence microscopy is constrained by the diffraction limit, while super resolution methods either require high intensity laser powers, suitable fluorophores, many of which have considerable toxicity, or heavy image post processing (*1*). Scanning electron microscopy (SEM) and correlated light and electron microscopy (CLEM) require fixation or introduce substantial disruption, making them unsuitable for live-cell imaging (*2*). Scanning ion conductance microscopy (SICM) has emerged as a powerful tool for non-contact, nanometre-resolution mapping of live-cell surfaces (*3*). By measuring ionic current changes through a nanopipette that scans over the cell surface, SICM generates quantitative topographical maps with nanometre precision, while avoiding mechanical interaction with the sample (*4*). This makes SICM particularly suitable for imaging delicate cell types such as neurons, epithelial cells, cardiomyocytes and even such complex tissue structures as glomerulus (*5–8*). Furthermore, SICM has enabled detailed studies of membrane protrusions, ion channel distribution, and cellular mechanosensitivity, demonstrating its immense potential for probing cell membrane morphology and dynamics (*9–11*).

Despite the above strengths, SICM is fundamentally limited to two-dimensional (2D) or quasi three-dimensional (3D) representations of the cell surface and does not provide access to volumetric information inside the cell. Yet, the cell membrane operates in close coordination with internal cellular processes, and many surface dynamics events are influenced by or contribute to changes within the cell. Comprehensive understanding of such events, therefore, requires simultaneous observation of membrane topography and intracellular activity. One promising approach is to combine SICM with volumetric fluorescence imaging, offering the ability to monitor intracellular events, especially those are proximate to plasma membrane (*12*). However, integrating these two modalities exhibits several challenges. Firstly, SICM achieves nanometre-scale resolution through high-precision piezo-controlled scanning, moving sample stage for volumetric fluorescence imaging can lead to spatial mismatch between the two systems, or significantly slow down acquisition speed due to the need for sequential, alternating movement. Secondly, moving the sample stage can also introduce vibrations that can compromise SICM stability and measurement accuracy. Thirdly, the physical setup of SICM typically places the nanopipette directly above the sample, leaving limited vertical space for integrating additional optical components. Overcoming these challenges will require a volumetric imaging strategy that minimises vibration while avoiding sample movement during acquisition.

Oblique plane microscopy (OPM) offers an ideal complementary approach to overcome the above challenges (*13*). Unlike conventional volumetric imaging methods that require sample stage movement, OPM illuminates the specimen with a thin, oblique light sheet generated from a single objective positioned beneath the sample. By rapidly scanning the light sheet through the cell using a galvo mirror, OPM achieves optical sectioning without physically moving the sample. The resulting fluorescence images can be computationally stacked to reconstruct a 3D image of intracellular structures. Without sample movement and using a single objective for both sample illumination and fluorescence detection render OPM inherently compatible with the scanning mechanism and space requirements of SICM. Moreover, the light-sheet-based illumination confines excitation to a narrow plane within the cell, significantly reducing phototoxicity and photobleaching, making OPM particularly suited for long-term live-cell imaging and dynamic studies, such as intracellular dynamics, mitotic events, and calcium signalling (*14–17*).

In this study, we present a novel implementation of a compact correlative SICM-OPM system designed for comprehensive live-cell imaging, utilising the complementary strengths of both modalities. We describe the optical and mechanical integration of the system, along with dedicated pipelines for cross-modality image registration and high-resolution point-based reconstruction. We demonstrate the versatility of the system across a range of live-cell imaging tasks, including transverse-tubule remodelling in live cardiomyocytes under drug treatment, and precision-guided nanopipette injection into neural cells and unicellular organisms such as diatoms. Importantly, our SICM-OPM system is fully compatible with standard inverted microscope frames, facilitating its integration into conventional imaging workflows. Overall, we aim to establish a versatile SICM-OPM platform for correlative imaging across structurally and functionally diverse biological systems.

## 2. Methods

### 2.1 OPM system description

Fig.1 shows the schematics of the OPM optical setup. A high NA, 100× silicone oil immersion objective (O1, NA 1.35, MRD73950, Nikon) is used both to provide an oblique, sheet-like illumination within the sample and to collect the fluorescence emission. The collimated fluorescence emitted from O1 is then focused by a 200 mm Nikon tube lens (TL1), housed within a Nikon Eclipse TE2000-U microscope frame. A pair of relay lenses (L7 and L8, AC254-100-A-ML, Thorlabs) is positioned immediately after the microscope camera port to extend the optical path. In the common path between L7 and L8, a galvanometric mirror (GM, GVS201, Thorlabs) is placed at the focal plane of L7 to enable light-sheet scanning during image acquisition. The fluorescence beam reflected from the galvo mirror then passes through a dichroic mirror (DM2, Di01-R406/488/561/635-25×36, Semrock), which redirects it upwards via a vertical periscope system (VP, RS99/M, Thorlabs) to the upper detection layer. On the upper layer, the beam is collimated by a second tube lens (TL2), which comprises two commercial achromatic doublets (ACT508-500-A-ML and ACT508-1000-A-ML, Thorlabs), selected using the Doublet Selector (*18*). An intermediate image is subsequently formed at the front focal plane of a secondary 40× air objective (O2, NA 0.95, CFI Plan Apochromat Lambda D, Nikon). This objective is mounted on a manual translation stage (LX30/M, Thorlabs), enabling precise pupil alignment. A glass-tipped tertiary objective (O3, NA 1.0, AMS-AGY v1.0, Calico Labs), also mounted on a manual translation stage (LX30/M, Thorlabs), is placed head-to-head with O2 at a 35° tilt. Finally, the image is focused onto a camera (sCMOS, PRIME-BSI-EXP, Photometrics) via a tube lens (TL3, TTL200-A, Thorlabs) and a motorised emission filter wheel (FW, FE103/M, Thorlabs). A critical aspect of the OPM alignment is ensuring that the back focal plane of O1, the galvo mirror (GM), and the back focal plane of O2 are optically conjugated (*19*). This alignment ensures that scanning the galvo mirror does not alter the beam or image position laterally at these planes, allowing accurate lateral scanning without introducing image distortion.

**Fig. 1.**
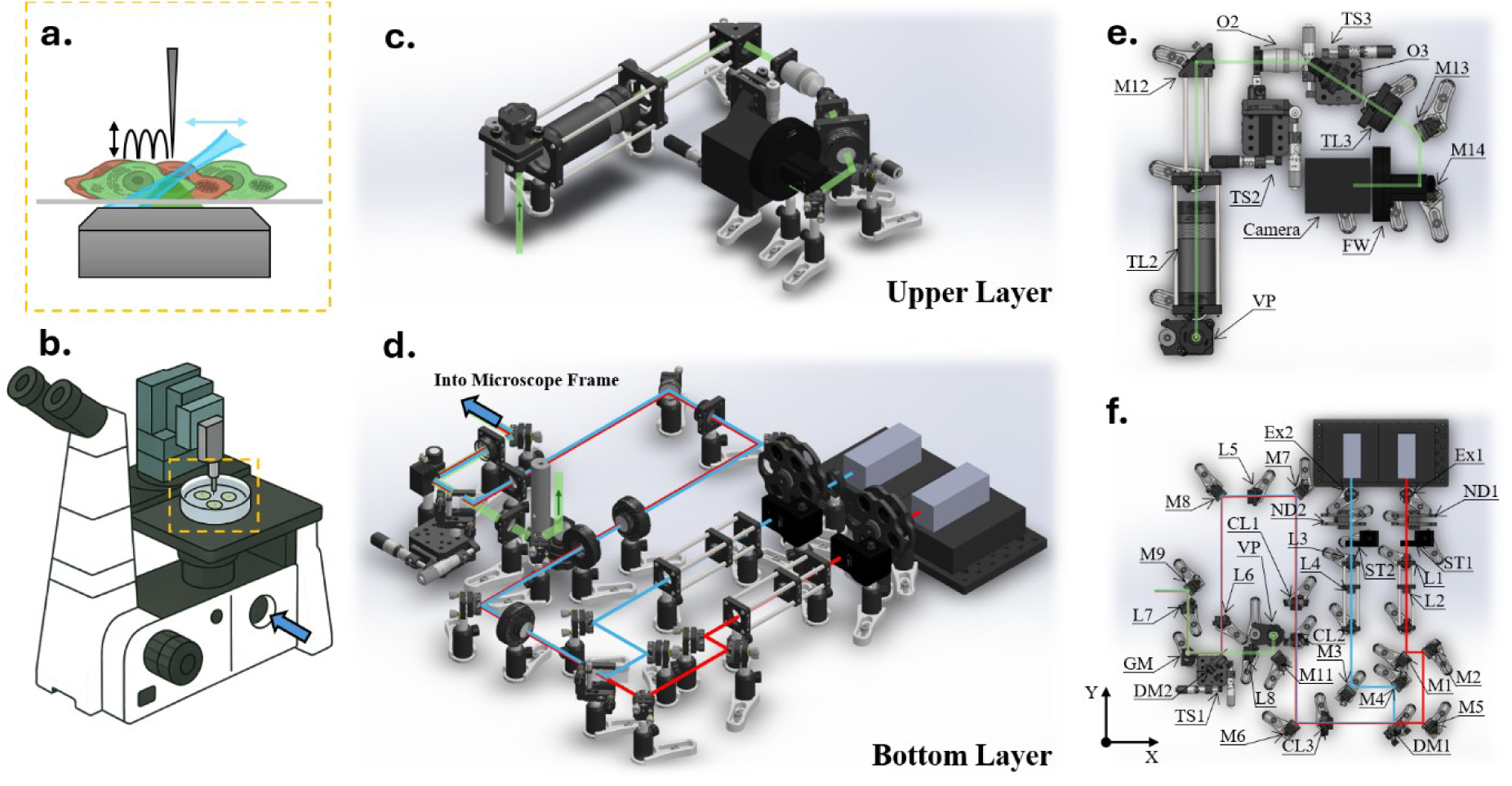
Layout of the SICM-OPM system. The OPM system comprises two distinct optical layers. (a) Schematic illustration of the SICM-OPM imaging principle. A hopping-mode SICM nanopipette (black arrow) performs non-contact nanoscale topographical imaging of the cell surface, while a tilted light sheet scans laterally (blue arrow) to enable fast volumetric fluorescence imaging. (b) Simplified diagram showing the SICM-OPM system integrated into a standard inverted microscope frame. The SICM head is mounted above the sample stage, while the OPM detection and illumination optics are coupled into the side ports of the microscope frame. (c, d) 3D perspective views of the upper and bottom layers, respectively, illustrating the spatial arrangement of system components. (e) Top-view schematic of the upper layer, showing the detection path. (f) Top-view schematic of the bottom layer, detailing the common path and detection path. O, objective; L, lens; M, mirror; TL, tube lens; FW, filter wheel; DM, dichroic mirror; CL, cylindrical lens; VP, vertical periscope; TS, translation stage; Ex, excitation filter; GM, galvo mirror; ST, optical beam shutter; ND, neutral density filter; EF, emission filter.

The illumination path is positioned on the bottom layer of the optical setup and begins with two laser sources (488 nm and 638 nm, 06-MLD-180/200, Cobolt, HÜBNER). Each laser is directed through an excitation filter (Ex1, FF01-637/7-25, Semrock; Ex2, MF475-35, Thorlabs) followed by a dedicated neutral density filter wheel (ND1 and ND2, FW2AND, Thorlabs) and a beam shutter (ST1 and ST2; SH05R/M, Thorlabs). The beams are then passed through individual beam expanders, each comprising a pair of identical achromatic doublets (L1–L4, AC127-025-A-ML, Thorlabs) with 1× magnification. The 488 nm beam is reflected by a dichroic mirror (DM1, Di02-R488-25×36, Semrock), while the 638 nm beam is transmitted through DM1, allowing the two beams to combine and overlap downstream. After beam combination, the light passes through a cylindrical lens (CL1, ACY254-100-A, Thorlabs), which focuses the beam along one axis while leaving the orthogonal axis unaffected. An achromatic doublet (L5, AC254-250-A-ML, Thorlabs) then collimates the focused axis and simultaneously begins to focus the previously unaffected direction. A second doublet (L6, AC254-100-A-ML, Thorlabs), paired with L7, further relays the beam. The beam is then directed onto a galvanometric mirror (GM) via a dichroic mirror (DM2). The beam is ultimately focused through tube lens TL1 and objective O1 onto the sample plane, forming a light sheet in one axis while remaining collimated in the orthogonal direction (see Fig. S1(a)). To maintain the light-sheet thickness while extending its width, a cylindrical lens pair (CL2, ACY254-200-A, Thorlabs; CL3, ACY254-050-A, Thorlabs) is introduced between CL1 and the dichroic mirror DM1. This lens pair expands the beam size by a factor of 4× in one direction, thereby increasing the light-sheet width without compromising its thickness. It is worth noting that all three cylindrical lenses (CL1, CL2, and CL3) are aligned with the same axis of curvature, ensuring that their focal power is applied consistently along the same direction to ensure uniform beam shaping across the optical path.

### 2.2 OPM imaging control

This OPM setup generates a measured thickness of approximately 1.7 µm and a full-width-at-half-maximum (FWHM) width of 45 µm at the sample plane. Lateral scanning was achieved using the galvo mirror (GM), controlled via a field programmable gate array (FPGA) board and custom LabVIEW programme. For precise control of the galvo mirror angle, two digital control ports on the galvo mirror controller are connected to the digital output channels (DO1 and DO2) on the FPGA board. During system alignment, the mirror was driven with a fixed voltage of 1.5 V (see Fig. S2(b)). During imaging mode, the galvo was scanned across 250 discrete steps with a voltage increment of 0.01 V per step, resulting in a lateral scan range of approximately 105 µm (see Fig. S1(b), (c)).

The LabVIEW program (see Fig. S2(a)) also includes manual controls for the galvo mirror, enabling convenient sample monitoring. The scanning centre can be digitally adjusted within the program (see Fig. S2(b)), avoiding the need for mechanical stage repositioning in some cases.

For synchronised galvo scanning and image acquisition, the camera was configured in the Edge Trigger mode via Micro-Manager 2.0. In this mode, each frame was triggered by the rising edge of an external trigger signal generated through an analogue output port (AP1) on the FPGA. The camera’s TRIG RDY output was connected to an analogue input port (AP2) on the FPGA, enabling the system to detect when the camera is ready to receive the next trigger. Once a high signal is detected on AP2, the LabVIEW programme will send a rising edge via AP1 to trigger the next frame.

Additional four analogue output ports (AP3–AP6) on the FPGA were used to control the emission filter wheel (FW) and two laser controllers, with two ports connected to the filter wheel’s analogue inputs and the other two to the laser control inputs.

In single-colour imaging mode, the emission filter wheel was fixed at a selected emission filter. In two-colour imaging mode, the filters were automatically switched after each volumetric scan, as specified in the LabVIEW control sequence. Lasers were modulated in analogue mode via LabVIEW and were only turned on during image acquisition to minimise photobleaching.

### 2.3 SICM description

The SICM scan head was mounted on the sample stage of the inverted Nikon Eclipse TE2000-U microscope frame (see Fig. 1(b). The coarse positioning of the SICM nanopipette was achieved by a PatchStar micromanipulator (Scientifica, UK), which provides 20 mm of travel in the X, Y, and Z axes.

Fine lateral (XY) scanning and positioning of the SICM pipette were performed using either an S-316.10 Piezo Z/Tip/Tilt Scanner (Physik Instrumente, Germany), offering a scan range of 98 × 98 µm, or an S23.ZT1S-C1 Tip/Tilt Scanner (CoreMorrow, China) with a 32 × 32 µm scan range. These Tip/Tilt scanners operate via three individually addressable piezo actuators arranged at the corners of an equilateral triangle, enabling precise control of both angular tilt and vertical displacement of the moving platform. To generate XY motion, the desired coordinates were converted into corresponding actuator voltages according to the *PZ 96E User Manual* (Physik Instrumente, Germany).

A custom Z-axis piezo actuator was assembled from a stack of 12 PICMA piezo rings (PD050.30, Physik Instrumente, Germany) and mounted on the moving platform of the Tip/Tilt scanner. This Z actuator offers a travel range of 22 µm and a resonant frequency of 4 kHz, as measured by an IDS3013 laser interferometer (Attocube, Germany). It is also equipped with a strain gauge sensor for closed-loop control. All piezo actuators and sensors were controlled and read out using a Piezo Control System (ICAPPIC Ltd., UK). The microscope system was operated using custom HPICMScanner software and the ICAPPIC Universal Controller (ICAPPIC Ltd., UK).

SICM nanopipettes were fabricated from borosilicate glass capillaries (outer diameter 1.0 mm, inner diameter 0.5 mm; B-100-50-10, World Precision Instruments) using a P-2000 laser puller (Sutter Instruments, USA). Nanopipettes used for topographical and mechanical imaging were pulled using a two-line program with the following parameters: Line 1: HEAT 350, FILAMENT 3, VELOCITY 30, DELAY 200, PULL 0 /Line 2: HEAT 340, FILAMENT 2, VELOCITY 27, DELAY 160, PULL 160. Note that these pulling parameters are specific to the individual instrument used and may require optimisation for different setups.

### 2.4 Imaging processing methods

The image processing in SICM-OPM involves SICM height map reconstruction (flowchart see Fig. S3(a)), OPM 3D image reconstruction (*20*), and correlative image registration (flowchart see Fig. S3(b)). For SICM, the raw topological data encoded as a 2D RGB heat map is first converted to a grayscale height map, which is further transformed into a 3D binary volume with a uniform voxel size along the X, Y, and Z axes. Within this volume, the value of a voxel (*X*, *Y*, *Z*) is determined by:

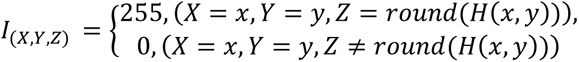

where *x*, *y*, and *H*(*x*, *y*) are the coordinates and height value of
the corresponding pixel in the 2D height map, respectively. This transformation effectively reconstructs the SICM topography as a binary shell within the 3D volume.

In parallel, the OPM 3D image reconstruction is performed using a custom Python script (*20*). The script first calls ImageJ to crop the region of interest (ROI) from the raw OPM image stack, ensuring that only the relevant portion of the data is processed. It then performs deskewing to reorient the tilted volume into its correct spatial alignment, making it suitable for quantitative analysis.

Next, the SICM binary shell within the 3D volume and the OPM 3D image volume are spatially aligned using a custom MATLAB-based image registration pipeline. Due to the higher lateral resolution of SICM, the raw OPM volume is first rescaled in the X and Y dimensions to match the pixel resolution of the SICM data. A 2D grayscale version of the SICM height map is first used as the alignment reference. The rescaled OPM volume was loaded as a 3D array, and a candidate search centre, defined by a user-input XY coordinate and a Z-range, was specified to constrain the registration search region. The registration is then performed by scanning through each Z slice of the rescaled OPM stack within the defined range and testing local XY shifts (ΔX, ΔY) within a specified radius. For each potential alignment, a similarity metric (2D correlation coefficient generated by MATLAB *corr2* function) is calculated between the SICM height map and the OPM slice. The best-matching position is identified based on the maximum similarity score. Once the optimal registration parameters were determined, the corresponding sub-volume of the OPM stack was extracted and saved as the aligned OPM image stack. This aligned volume was then rescaled in the Z direction to restore isotropic voxel dimensions for downstream analysis and visualisation.

To aid the registration process, a fluorescent spatial marker was introduced before each experiment by filling the SICM nanopipette with fluorescently labelled dextran (D22914, Thermo Fisher). The labelled pipette was imaged using the OPM, providing a visible reference point in the fluorescence volume that corresponded to the SICM scanning centre. This marker ensured that the initial search region could be centred near the true SICM imaging location, improving both accuracy and efficiency.

### 2.5 High-resolution point-based (HRPB) reconstruction of T-tubules from OPM fluorescence images

To improve the visualisation of T-tubule structures in cardiomyocytes, a custom MATLAB-based image reconstruction pipeline was developed to generate high-resolution point-localisation image stacks from raw OPM fluorescence images.

Each frame of the OPM stack was first converted to double precision and processed using a Laplacian of Gaussian (LoG) filter to enhance and detect point-like features. The LoG filter parameters were chosen based on the measured system point spread function (0.46 µm FWHM). A fixed intensity threshold was applied to suppress background fluorescence, and local maxima in the filtered image were identified as candidate T-tubule localisations.

To isolate meaningful structures and eliminate background artefacts, a binary cell mask was generated based on intensity thresholding. Small objects and features near image borders were removed, and localisation candidates falling outside the cell mask were excluded from further analysis.

For each accepted localisation, the local signal intensity was quantified using a circular region of interest. Alternatively, an option was provided to assign a uniform intensity to all localisations to better visualise the geometric organisation of T-tubules, independent of fluorescence variation.

The coordinates of detected localisations were then used to construct a high-resolution image by upsampling the original frame by a factor matched to the pixel resolution of the corresponding SICM data. Each localisation was rendered as a Gaussian spot, using a kernel whose width was scaled to reflect the measured system PSF in the upsampled space, see Section 3.1. A high-resolution image was generated for each frame by summing the contributions of all localisations (see Fig. S4).

While the current implementation does not perform super-resolution reconstruction beyond the diffraction limit, it enhances the spatial representation of subcellular features by resampling and re-rendering localised signals on a finer spatial grid. This approach improves structural clarity and supports quantitative analysis of T-tubule architecture, without requiring hardware-based super-resolution methods (see Section 3.2). Notably, the underlying framework allows for future extension to true super-resolution reconstruction, by adjusting the LoG filter size to match a smaller PSF corresponding to a higher resolution target.

### 2.6 SICM injection

Fluorescently labelled cargos were delivered into target cells—including *Coscinodiscus radiatus* diatoms and neuroblastoma cells—using a localised electroporation approach enabled by SICM nanopipette injection. The nanopipette, pre-loaded with the appropriate cargo solution, was precisely positioned vertically above the target cell surface. The pipette was advanced toward the cell surface until a 0.5% drop in ion current was detected relative to the baseline (i.e., when the pipette was distant from the surface), indicating proximity to the cell. The pipette was then lowered an additional 2 µm to ensure membrane contact.

A sequence of 20 electroporation pulses was delivered by applying a voltage protocol to the reference electrode in the bath: −10 V for 50 ms, +10 V for 5 ms, and −0.1 V for 2 ms per episode. Nanopipettes for this injection experiment were fabricated using a P-2000 laser puller (Sutter Instruments, USA) with the following two-line program: Line 1: HEAT 280, FILAMENT 3, VELOCITY 20, DELAY 150, PULL 0 /Line 2: HEAT 300, FILAMENT 4, VELOCITY 15, DELAY 120, PULL 100. Note: Pulling parameters are specific to the individual puller used and may require optimisation for other instruments.

### 2.7 Biological sample preparation methods

#### 2.7.1 Diatom cultivation

The diatom *Coscinodiscus radiatus* (CCMP312) was obtained from the Provasoli-Guillard National Center for Marine Algae and Microbiota (NCMA, USA). The culture was grown in artificial seawater medium (*21*) and in a controlled environmental growth room at 23 °C with an illumination of 50 μmol·m-2·s-1 (12 h light / 12 h dark) at Silwood park campus of Imperial College London. The seawater medium was prepared using aseptic techniques and laboratory materials were acid-cleaned. Chemicals of ACS grade or higher purity were used, and the artificial seawater was 0.2-µm filtered (polycarbonate filters, Merck Millipore Ltd.) and kept at 4 °C in dark before use. Fresh batches were inoculated in a microbiological safety cabinet by transferring 5 mL of culture to 35 mL of media in a new 40 mL culture flask fortnightly.

#### 2.7.2 Adult rat cardiomyocytes

Adult rat ventricular cardiomyocytes (ARVMs) were isolated from male adult Sprague–Dawley (SD) rats (150–250 g) using a Langendorff perfusion system. Following isolation, approximately 10,000 cardiomyocytes were seeded onto 35 mm glass-bottom dishes (P35G-1.5-14-C, MatTek) pre-coated with laminin. Cells were initially incubated in Minimum Essential Medium (MEM, Cat.#31095029, Gibco™) supplemented with 10% fetal bovine serum (FBS, Thermo Fisher) and 1% Antibiotic–Antimycotic Solution (Cat.#A5955, Sigma) for 1 hour at 37 °C in a humidified atmosphere with 5% CO2. The medium was replaced with serum-free MEM for subsequent treatments.

To induce T-tubule disruption, cells were treated with 300 µM of imipramine hydrochloride (Cat.#7841, Tocris Bioscience) diluted in serum-free MEM. The ARVMs were incubated with the imipramine solution for 15 minutes at 37 °C, 5% CO2.

For membrane staining, a Di-8-ANEPPS working solution was prepared by diluting a 2 mM stock to a final concentration of 20 µM in a high-potassium buffer (120 mM K-gluconate, 25 mM KCl, 2 mM MgCl₂, 1 mM CaCl₂, 2 mM EGTA, 10 mM HEPES, 10 mM glucose, adjusted to pH 7.4). The solution was sonicated in a water bath at 55 °C for 5 minutes.

Cells were washed twice with high-potassium buffer and stained with the Di-8-ANEPPS solution for 5 minutes at room temperature (RT) in the dark, followed by high-potassium buffer washing twice and immediate imaging.

#### 2.7.3 SH-SY5Y human neuroblastoma cell line

SH-SY5Y human neuroblastoma cells were cultured at 37 °C and 5% carbon dioxide in Roswell Park Memorial Institute medium (RPMI 1640, Thermo Fisher) with 10% fetal bovine serum (FBS, Thermo Fisher). Trypsin-EDTA (0.25%, Thermo Fisher) was used to detach cells from the culture plate, and culture medium was refreshed every 2-3 days. SH-SY5Y cells were harvested at ∼70% confluency and subsequently plated on glass-bottomed dishes (P35G-1.5-14-C, MatTek). MatTek dishes were coated with Poly-D-lysine (0.1 mg/mL) and collagen from rat (0.1 mg/mL) and incubated at RT for 1 h, prior to washing three times with Dulbecco’s phosphate-buffered saline (DPBS) and subsequent cell seeding.

#### 2.7.4 *α*-synuclein monomers tagged with Alexa Fluor 488

α-synuclein carrying the A140C mutation (A140C α-syn) was purified using a previously reported protocol (*22*). Briefly, the protein was overexpressed in *E. coli* BL21(DE3) overnight with 1 mM IPTG, followed by cell lysis, streptomycin sulfate precipitation of nucleic acids, and ammonium sulfate precipitation of proteins. α-Syn was further purified by ion-exchange chromatography on a HiPrep Q HP 16/10 column (Cytiva) using a 0–1 M NaCl linear gradient, followed by size-exclusion chromatography on a HiLoad 16/600 Superdex 75 pg column (GE Healthcare) in standard PBS. All steps were performed in the presence of 0.5 mM DTT.

Labelled α-syn monomers were obtained by conjugation of A140C α-syn with Alexa Fluor 488 C5-maleimide (Thermo Fisher). A140C α-syn was buffer-exchanged using Zeba Spin Desalting Columns (7K molecular weight cutoff) (Thermo Fisher) to remove the DTT from the storage solution. The reaction was carried out according to the manufacturer’s instructions.

## 3. Results

### 3.1 Optical system characterisation with fluorescent beads

We first estimated the theoretical resolutions of the system based on the following parameters: an effective numerical aperture NA (35° tilted configuration, see Fig. 1(e)) of 1.33, emission wavelength of 525 nm, bead diameter of 100 nm, camera pixel size of 6.5 μm, and an effective lateral magnification of 56×. Assuming that these contributions to the point spread function (PSF) can be approximated as independent Gaussian distributions, the theoretical lateral FWHM was estimated to be 0.41 μm, and the theoretical axial FWHM was 0.73 μm.

To experimentally characterise the system resolution, we imaged sub-diffraction fluorescent beads (TetraSpeck Microspheres, T7279, 100 nm nominal diameter, Thermo Fisher) embedded in 1.2% agarose. A three-dimensional maximum intensity projection of the bead sample is shown in Fig. 2(a). Since the OPM system is configured to image the upper surface of cells, the tertiary objective O3 was adjusted axially to focus approximately 10 μm above the coverslip. The optimal focus region—defined by the thickness of the light sheet (∼1.7 µm)—was approximately 7 μm in depth. Therefore, only beads located within this range were analysed to ensure accuracy and consistency.

**Fig. 2.**
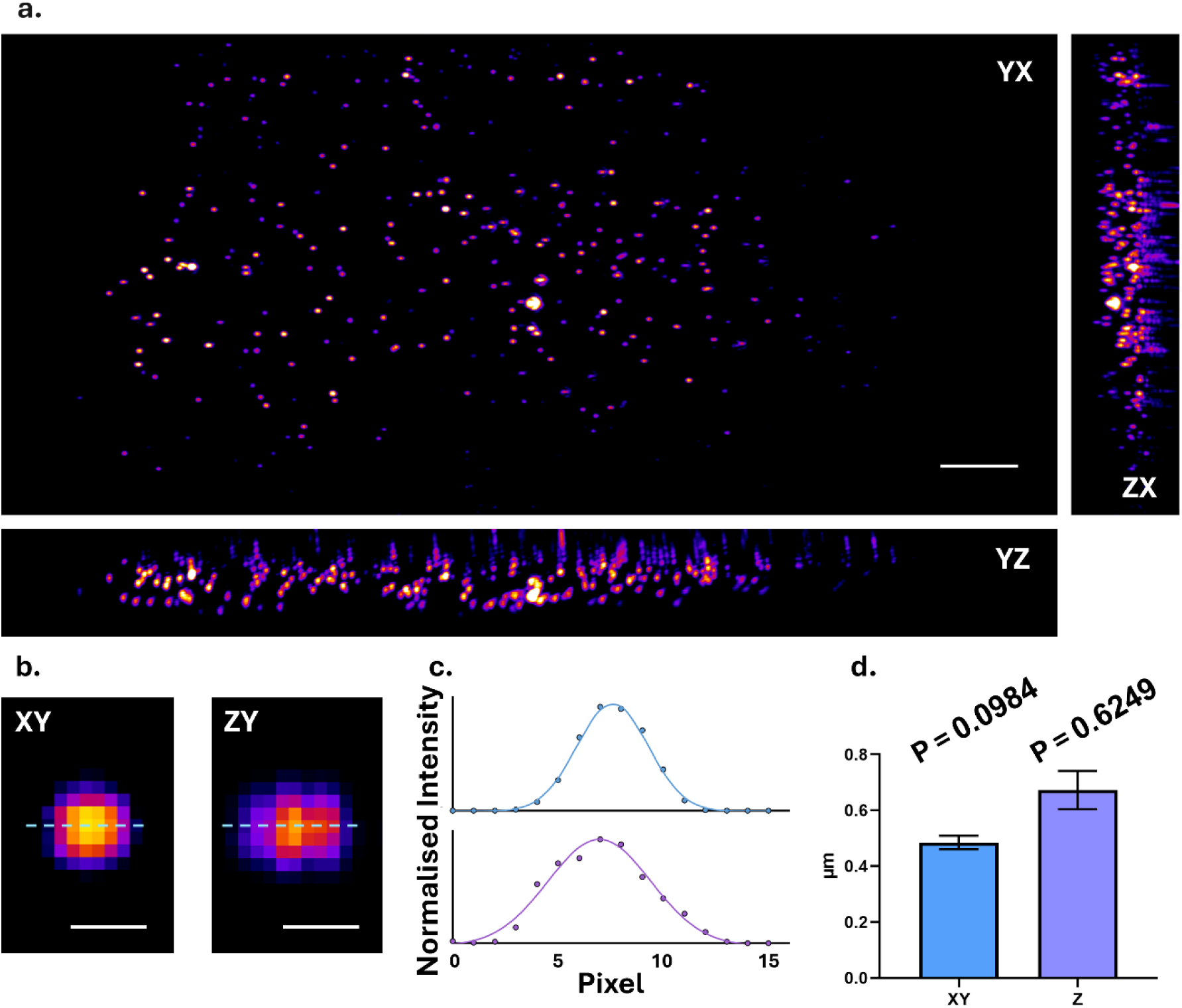
Optical characterisation of the OPM system using 100 nm diameter fluorescent beads. (a) Maximum intensity projections (YX, YZ, and ZX) of a 3D bead stack imaged using the OPM system. Scale bar: 10 µm. (b) Example XY and ZY views of a single fluorescent bead. Scale bar: 1 µm. (c) Normalised intensity profiles (dots) and corresponding Gaussian fits (curves) extracted along the dashed lines in panel (b) for the example bead. The measured lateral and axial FWHM were 0.46 μm and 0.67 μm, respectively. (d) Quantification of lateral and axial resolution based on FWHM measurements of 90 individual beads. Data are presented as mean ± standard deviation. All data groups passed the D’Agostino-Pearson normality test, confirming Gaussian distribution of measured FWHM values (P > significance level α = 0.05).

The lateral resolution was then assessed by measuring the FWHM of the intensity profile across the bead image in the XY plane. The axial resolution was determined from the FWHM of the Z-axis intensity profile extracted through the brightest pixel of each bead (see Fig. 2(b), (c) for an example).

Averaging across 90 fluorescent beads, yielding mean FWHM values of 0.48 ± 0.02 μm (lateral) and 0.67 ± 0.07 μm (axial) (Fig. 2(d)). These experimental values are in good agreement with theoretical estimates, with slight deviations attributable to minor degradation in image quality from remote refocusing, since imaging was taken place above the primary focal plane of O1 (*19*). Notably, the distributions of both lateral and axial resolution measurements passed the D’Agostino–Pearson test for normality, supporting the robustness of the system’s resolution characterisation.

### 3.2 Correlative volumetric imaging of cardiomyocyte detubulation

Transverse tubules (T-tubules) are essential membrane structures in cardiomyocytes, formed by deep invaginations of the sarcolemma that extend into the cell interior (*23*). T-tubules enable the rapid and uniform transmission of action potentials throughout the cell and are essential for the excitation–contraction coupling in cardiomyocytes. Structural remodelling of T-tubules, alongside the loss of membrane z-grooves where T-tubule openings typically localise, is a hallmark of myocardial infarction and other cardiac pathologies (*24*). Importantly, the extent and pattern of T-tubule disruption vary across different disease models and may offer mechanistic insights into disease progression. For example, formamide-induced detubulation of rat cardiomyocytes occurs via osmotic shock and results in substantial loss of both surface z-grooves and internal T-tubule structures (*25, 26*). In contrast, cationic amphiphilic drugs (CADs), such as imipramine, have been reported to disrupt T-tubules by dissociating of the membrane scaffolding protein BIN1 from PI(4,5)P2 (*27, 28*), although direct 3D visualisation of this process has not been previously demonstrated.

To address this, we used the SICM-OPM platform to investigate imipramine-induced detubulation in adult rat ventricular myocytes (ARVMs) as a proof-of-concept demonstration of the system capability to simultaneously resolve distinct membrane features and intracellular structures. Fluorescence-labelled T-tubule structures (via Di-8-ANEPPS) and high-resolution surface topography were simultaneously acquired from the same region of interest (Fig. 3(a), Fig. S5). To enhance the visualisation of submembrane T-tubule networks, high-resolution point-based (HRPB) reconstructions (see Section 2.5) were performed on the OPM fluorescence data (Fig. 3(b)). Correlative SICM-OPM imaging showed that untreated ARVMs exhibited intact surface morphology and continuous, well-defined T-tubule structures (Fig. 3(c)-(e)). In contrast, ARVMs treated with 300 μM imipramine showed no apparent changes in surface z-groove topography in SICM images (Fig. 3(f)-(h), Fig. S6(g), (h)). However, SICM topography also revealed a significant enlargement of T-tubule openings (Fig. S6(i), with an average width increased of 35.5% ± 8.3% (n = 43), suggesting disruption of T-tubule integrity (Fig. S6(i)). Meanwhile, OPM fluorescence imaging indicated a substantial reduction in T-tubule density by 85.1% ± 6.7% (n = 10), and a decrease in T-tubule regularity by 65.9% ± 14.2% (n = 10), as evidenced by fragmented or absent T-tubules (Fig. 3(i), (j), Fig. S6(e), (f)). Together, these findings suggest that imipramine selectively impairs the internal T-tubule architecture while largely preserving surface morphology.

**Fig. 3.**
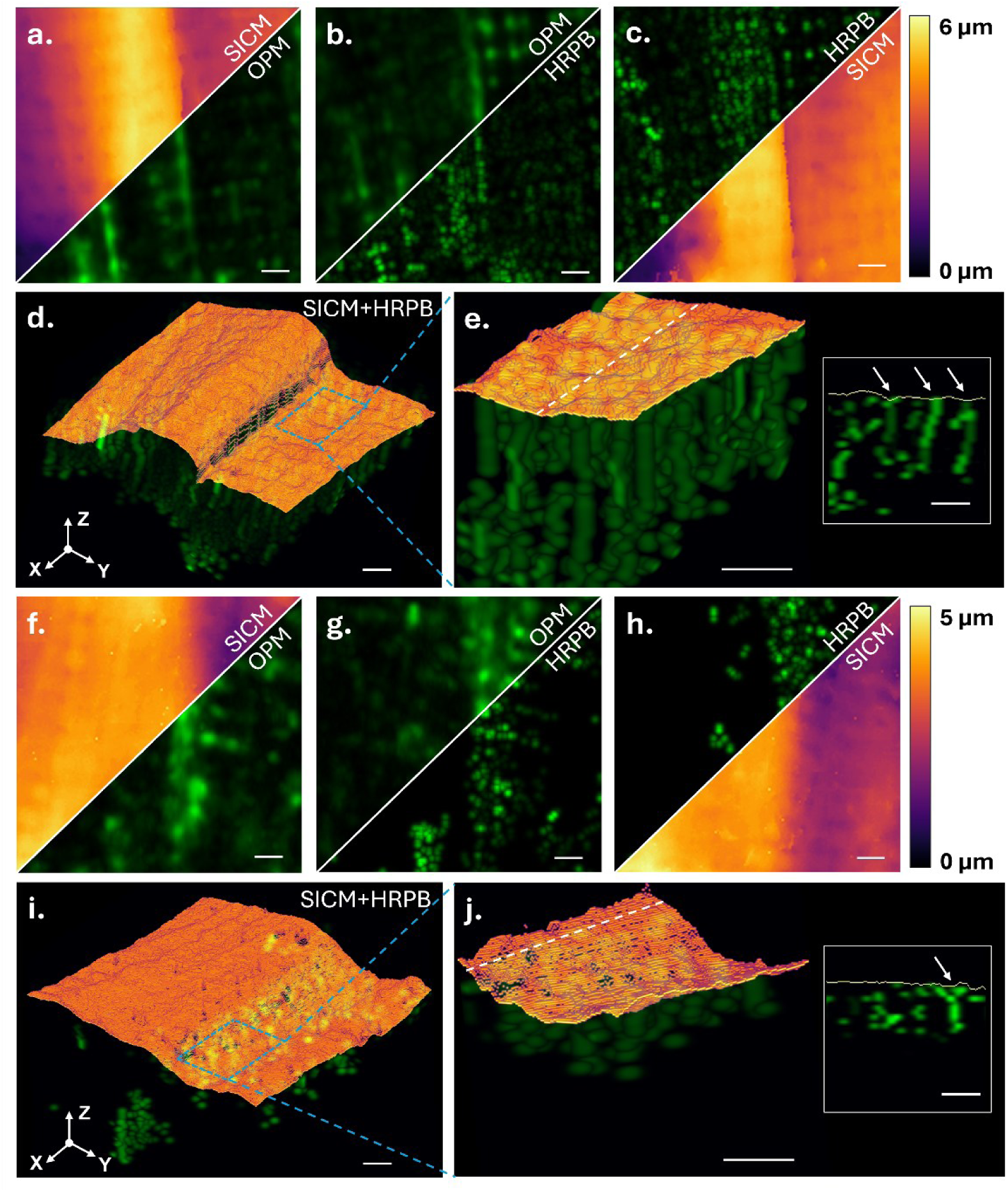
Correlative SICM-OPM imaging reveals distinct patterns of cardiomyocyte detubulation following imipramine treatment. (a-c) Correlative imaging (20 × 20 µm) of a control cardiomyocyte showing (a) SICM topography, (b) OPM fluorescence image, and (c) high-resolution point-based (HRPB) reconstruction of Di-8-ANEPPS-labelled T-tubules. (d) 3D rendering of the SICM+HRPB data from the same cell as (a–c), illustrating intact surface z-grooves and well-organised submembrane T-tubule structures. (e) Zoom-in of the region outlined in (d), highlighting the continuity of subsarcolemmal invaginations. Inset shows a cross-sectional intensity profile along the white dashed line. The white arrows mark clearly visible T-tubule openings. (f–h) Correlative imaging (20 × 20 µm) of a cardiomyocyte treated with 300 µM imipramine showing (f) SICM topography, (g) OPM fluorescence, and (h) HRPB reconstruction. (i) 3D rendering of SICM and HRPB data from the same treated cell, revealing preserved surface topography but disrupted internal T-tubule architecture. (j) Zoom-in of the region outlined in (i), where the T-tubule openings (white arrows) are markedly reduced compared to control. Inset: cross-sectional intensity profile showing loss of invagination continuity. Scale bar: 2 µm.

Importantly, the fluorescence imaging results from our SICM-OPM platform are highly consistent with previously confocal microscopy studies. In our prior work (*29*), ARVMs treated with 300 μM imipramine and imaged by confocal microscopy exhibited a similar reduction in T-tubule density and regularity, consistent with the values reported here. Comparable findings have also been reported by others under equivalent treatment conditions (*28*). These concordant findings not only attest to the robustness and accuracy of our SICM-OPM measurements but also highlight its unique advantage in capturing high-resolution 3D images of live cardiomyocyte T-tubule networks.

To our knowledge, this is the first demonstration of correlative imaging that simultaneously resolves both the surface topography and intracellular T-tubule architecture within the same live cardiomyocyte. Further studies will leverage this platform to investigate the dynamics and mechanisms underlying cardiomyocyte detubulation in greater depth. Overall, these results highlight the power of SICM-OPM for visualising membrane-associated ultrastructures and detecting subcellular pathological changes with high specificity.

### 3.3 Precision-guided nanopipette injection with SICM-OPM

Observing single-cell responses to the localised delivery of exogenous cargos offers valuable insights into a range of fundamental biological processes, including antigen recognition, inflammatory signalling, viral entry, and targeted drug delivery. The SICM-OPM platform facilitates such investigations by leveraging the nanopipette not only as a topographic probe, but also as a highly precise tool for electroporation and molecular delivery. Here, we demonstrate the system’s capability for high-precision, targeted single-cell injection in both *Coscinodiscus radiatus* diatoms and mammalian neuroblastoma (SH-SY5Y) cells, highlighting its versatility across phylogenetically and structurally distinct cell types.

We first performed targeted injection of FITC-Dextran (70 kDa, 46945, Sigma-Aldrich) into the girdle region of a *Coscinodiscus radiatus* diatom cell undergoing cytokinesis (Fig. 4(a), (b)). While diatom cells are enclosed within two rigid silica-based frustules, the girdle region remains mechanically softer and more accessible to nanopipette penetration, particularly during cell division before the completion of silica deposition (*30*). Fluorescence signals were predominantly localised inside the frustule space between the two closely apposed daughter cells, rather than within the cytoplasm. This region likely corresponds to the developing cleavage plane between two nascent valves, representing a transient and structurally delicate compartment during cytokinesis. The successful delivery of FITC-Dextran into this intermediate zone highlights the capability of the SICM-OPM platform in accessing shielded or mechanically constrained subcellular domains and potentially offers a new approach for gene delivery into diatoms.

**Fig. 4.**
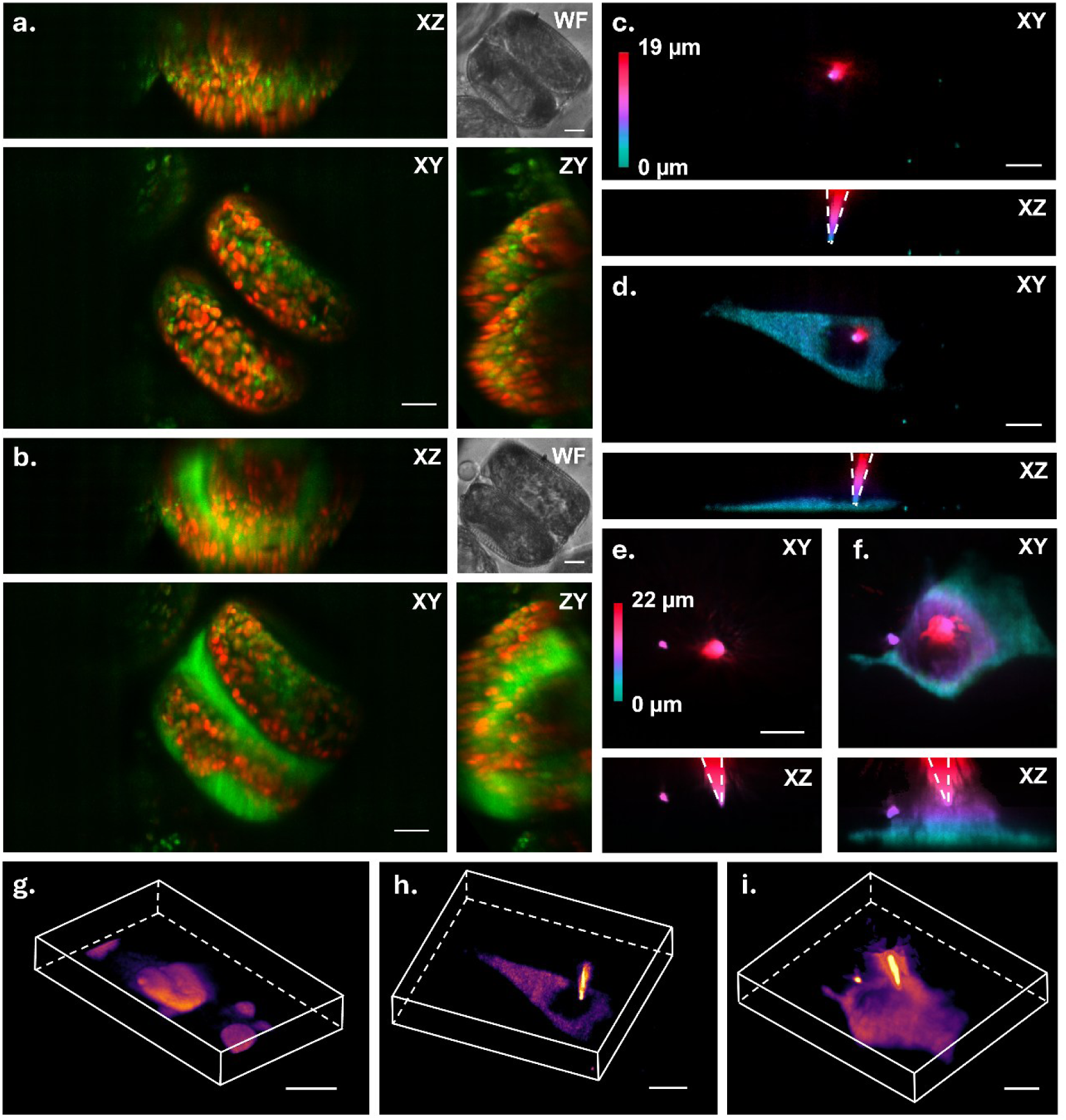
SICM-OPM enables volumetric visualisation of localised single-cell injection in diatoms and mammalian cells. (a, b) Maximum intensity projections (MIPs) of *Coscinodiscus radiatus* (CCMP312) before (a) and after (b) local electroporation-based injection of FITC-Dextran (70 kDa, 46945, Sigma-Aldrich). Each panel includes XY, XZ, and ZY views from OPM, along with a corresponding widefield (WF) image. (c, d) XY and XZ projections of an SH-SY5Y human neuroblastoma cell injected with fluorescein (5 μM, 46955, Fluka) into the cytoplasm. The SICM nanopipette position is indicated by white dashed lines. (e, f) An SH-SY5Y cell injected with 70 kDa FITC-Dextran before (e) and after (f) injection. Dashed lines mark the nanopipette location. (g) 3D rendering of an SH-SY5Y cell injected with fluorescein into the nucleus. (h, i) 3D views of the cells shown in (d) and (f), respectively, illustrating the spatial distribution of the injected material. All scale bars: 10 μm.

Next we applied the SICM-OPM system for intracellular injection of both low- and high-molecular-weight fluorescent reagents into SH-SY5Y cells and monitored their subcellular distribution. Cytoplasmic injection of fluorescein (5 μM, 46955, Fluka) resulted in strong intracellular fluorescence confined within the cytoplasmic volume but not in the nucleus (Fig. 4(c), (d), (h)). Despite the thin cytoplasmic layer in the cell, electroporation-based injection was achieved with high precision, highlighting the system’s fine positional control. In a separate experiment, fluorescein was selectively injected into the nucleus of another SH-SY5Y cell (Fig. 4(g)), further demonstrating subcellular targeting precision. Similar localisation patterns were observed upon injection of FITC-Dextran (70 kDa) (Fig. 4(e), (f), (i)), confirming the system’s compatibility with a range of molecular cargos.

To further demonstrate the multifunctionality of the SICM-OPM platform, we performed a proof-of-concept experiment combining targeted protein delivery with correlative topographical and mechanical mapping. Fluorescently labelled α-synuclein monomer was injected into an SH-SY5Y neuroblastoma cell (C1), with an adjacent non-injected cell (C2) serving as an internal control. OPM fluorescence imaging revealed an increase in fluorescence within C1 post-injection (Fig. 5(a), (b)), while C2 remained largely unchanged. Interestingly, a small fluorescent punctum emerged in C2 (highlighted by a red arrow), possibly indicating intercellular diffusion or endocytosis of α-synuclein. Notably, an autofluorescent feature visible in C1 prior to injection remained stable, confirming cell identification across time points.

**Fig. 5.**
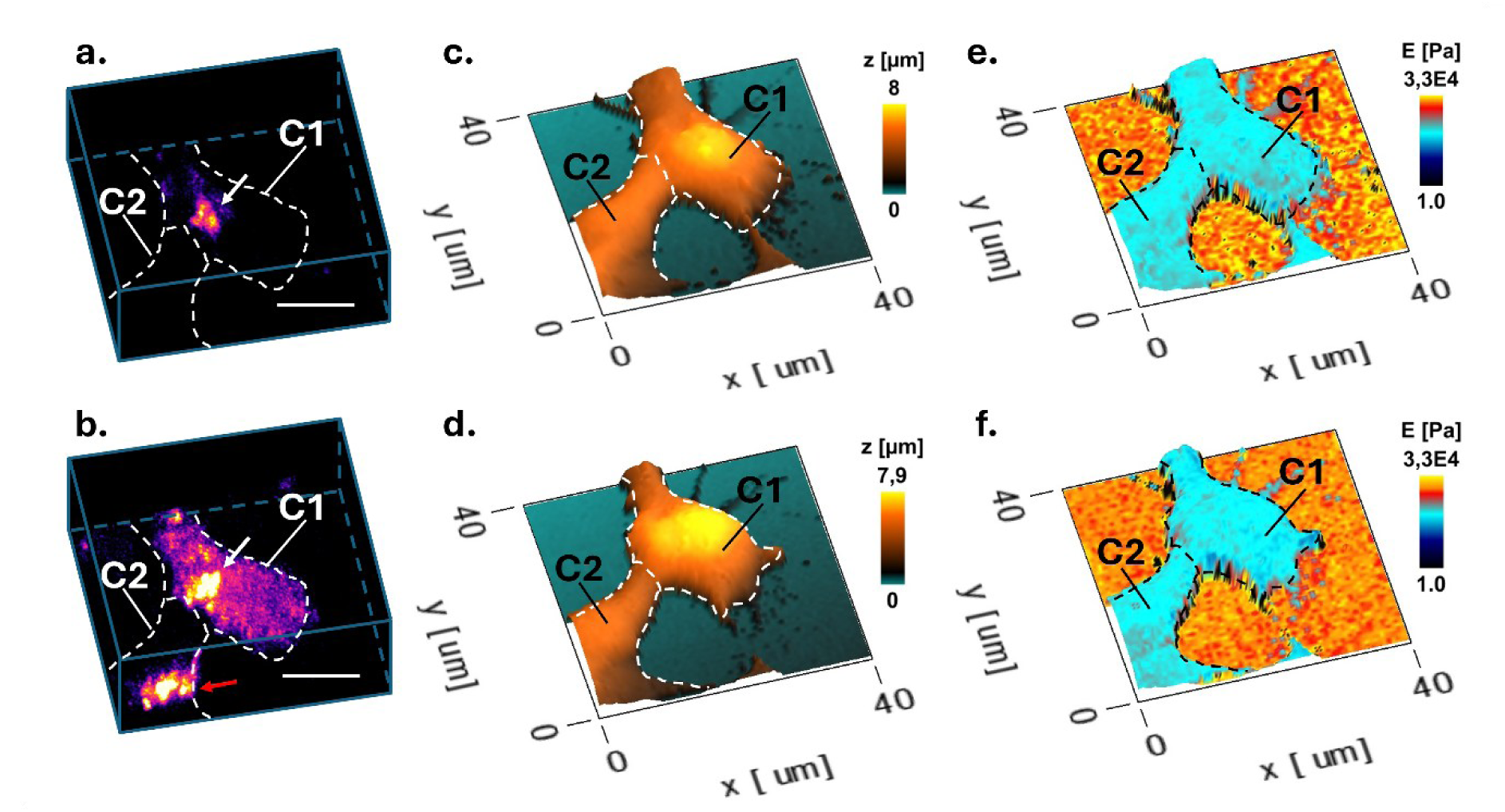
SICM-OPM enables single-cell α-synuclein delivery and mechanical mapping changes in injected and uninjected SH-SY5Y cells. (a, b) 3D OPM fluorescence images of two adjacent SH-SY5Y neuroblastoma cells (C1 and C2) before (a) and after (b) nanopipette-based electroporation of fluorescently labelled α-synuclein into C1. A persistent autofluorescent feature is visible in C1 pre- and post-injection (white arrow), while a new fluorescent punctum appears in C2 post-injection (red arrow), suggesting possible intercellular transfer. (c, d) Corresponding SICM topography maps acquired immediately (c) and 3 hours (e) after injection, showing no major morphological changes. (e, f) Young’s modulus (E) maps showing spatial stiffness distribution of the same field, highlighting time-dependent softening in the injected cell (C1) compared to the uninjected neighbour (C2). Cell boundaries are outlined with dashed lines. Colour bars: height (c, d), stiffness in pascals (e, f).

SICM-based topographical and mechanical imaging was conducted immediately after injection and again three hours later. 3D topographical scans showed no significant morphological changes over time (Fig. 5(c), (d)), whereas corresponding mechanical stiffness maps revealed changes in cellular biomechanics (Fig. 5(e), (f)). Immediately post-injection, C1 and C2 regions (Fig. S7(a)) exhibited comparable stiffness levels (Young’s modulus: C1, 446.0 ± 157.6 Pa; C2, 473.6 ± 150.2 Pa). Three hours post-injection, both cells exhibited softening, but the effect was more pronounced in the injected cell C1 (C1: 325.6 ± 103.4 Pa; C2: 439.4 ± 142.7 Pa; Fig. S7(b)). These results demonstrate the platform’s capability to deliver protein cargo with subcellular precision and to monitor subsequent mechanical changes in a time-resolved manner. Together, these findings present the broad utility of the SICM-OPM platform in delivering molecules with subcellular precision, while simultaneously capturing correlative topographical and mechanical responses. This integrated approach offers a powerful platform for investigating intracellular transport, cytoskeletal remodelling, and pathophysiological processes—such as protein aggregation in neurodegenerative diseases—with broad potential applications in neuroscience, immunology, and nanomedicine.

## 4. Discussion and Conclusions

The SICM-OPM system presents a highly versatile platform for precise single-cell manipulation and correlative imaging. In this study, we demonstrated its capabilities through direct visualisation of detubulation in rat ARVMs and precise delivery of fluorescent cargos with varying molecular weights into both diatom and mammalian cells. These examples rely on the dual functionality of the nanopipette, which can switch seamlessly between high-resolution topographic mapping and targeted intracellular delivery, expanding the system’s applicability to a broad range of biological investigations.

SICM has been combined with confocal microscopy in previous studies to enable simultaneous topographical and molecular visualisation (*12*), but these implementations faced significant technical limitations. In the earlier system, the scanning strategy required physical translation of the sample using a motorised stage to generate topographical maps. This approach not only introduced mechanical instability and motion artifacts but also severely constrained acquisition speed, rendering it suboptimal for dynamic live-cell imaging. Furthermore, confocal fluorescence imaging necessitates serial z-stack acquisition to match the 3D scanning volume of SICM, resulting in long imaging times, increased photobleaching, and phototoxicity. In contrast, SCIM-OPM addresses both challenges through key architectural innovations. Firstly, the sample remains stationary while the SICM nanopipette is precisely scanned using a piezoelectric micromanipulator, significantly reducing mechanical disturbance and enabling higher spatial and temporal stability. Secondly, the integration of OPM allows for instantaneous volumetric fluorescence imaging without the need for z-stack acquisition. This capability enables rapid 3D imaging over volumes comparable to those scanned by SICM, achieving true volumetric correlation within seconds.

The ability of SCIM-OPM to perform high-speed, high-resolution, and minimally invasive correlative imaging provides a substantial advantage for studying live cellular dynamics in 3D. One promising application is the study of mechanosensory neurons (*31*), which convert mechanical deformation into electrical or biochemical signals. SICM enables non-invasive, high-resolution probing of local membrane mechanics or specialised nerve terminals, while OPM can simultaneously capture intracellular calcium transients or neuronal firing using calcium indicators, thereby investigating stimulus-response coupling in mechanotransduction (*32*). Another potential application lies in the investigation of ferroptosis, a non-canonical form of programmed cell death driven by membrane lipid peroxidation (*33*). While the primary biochemical changes occur intracellularly—such as iron overload, reactive oxygen species (ROS) accumulation, and mitochondrial dysfunction—these processes also lead to secondary alterations in the plasma membrane’s nanomechanical properties and integrity. SICM can sensitively detect such nanoscale disruptions on the cell surface, including changes in stiffness, which reflect ferroptotic damage. Concurrently, OPM can image the intracellular biochemical events using appropriate fluorescent reporters (*34*), enabling a comprehensive, correlative view of the spatiotemporal progression of ferroptosis. The targeted delivery capability of the SICM-OPM system further extends its utility to functional studies of exogenous gene and protein expression. This is particularly promising in the context of genetic engineering of diatoms where reliable and minimally invasive delivery methods are much required (*35*). In addition, plasmids can be precisely injected by the nanopipette into the nucleus or cytoplasm to induce gene expression, while cellular dynamics are monitored in real time by OPM. Similarly, direct delivery of functional proteins, such as transcription factors, signalling molecules, cytokines, or even protein aggregates, enables investigation of their immediate effects on cell phenotype and signalling pathways. These capabilities can be readily integrated with downstream single-cell omics technologies, such as single-cell RNA-seq or ChIP-seq, to correlate functional responses with transcriptional or epigenetic changes, providing a comprehensive, multi-layered view of cell state. This feature opens up new possibilities for studying compartment-specific signalling, organelle-specific drug responses, or spatially restricted biomolecular interactions. Such targeted perturbations are valuable for drug target validation, enabling direct investigation of target localisation, function, and the phenotypic consequences of local manipulation.

Furthermore, several future enhancements could significantly extend the system’s versatility. Firstly, optimising the light-sheet-based OPM for a larger imaging volume would enable comprehensive volumetric imaging of tissue and organoids. Secondly, by performing motorised stage translation between sequential volumetric imaging sessions, the system could achieve high-resolution imaging across extended or large-format biological specimens. Thirdly, the integration of an environmental chamber or live-cell incubator would support long-term physiological experiments under tightly controlled conditions—crucial for studies on development, regeneration, or pharmacological response. On the SICM side, expanding the lateral scanning range and improving the responsiveness of piezo actuators would allow for faster, large-area imaging without compromising resolution, paving the way for higher-throughput investigations.

Overall, the SICM-OPM platform represents a powerful tool for live-cell investigations, offering unique capabilities for membrane morphology characterisation, targeted delivery, and functional imaging in a label-compatible, minimally invasive manner. These integrated capabilities are expected to empower fundamental discoveries across diverse biological fields, including neurodegeneration, inflammation, immune response, and cancer biology.

## 5. Funding and Acknowledgements

WH, ZZ, DK and AS acknowledge support from EPSRC (EP/W012219/1, EP/W015005/1, EP/X034968/1). AL acknowledges support from Imperial College London and the China Scholarship Council (CSC) Scholarship (File No. 202309370006). YW acknowledges support from Early Career Pathway Award of the CRUK International Alliance for Cancer Early Detection (EDDAPA-2024/100006). DK acknowledges support from the Royal Society. JG acknowledges support from British Heart Foundation (RG/F/22/110081). DMV acknowledges support from Alzheimer’s Society (Dementia Research Leader Fellowship AS-DRL-24-012). FAA acknowledge support from UKRI Future Leaders Fellowship (MR/S033947/1, MR/Y003616/1) and Alzheimer’s Research UK (Major Project Grant ARUK-PG2019B-020). FL acknowledges support from UK Natural Environment Research Council Grant (NE/V01451X/2). The authors thank the Centre of Excellence Cellular Mechanosensing and Functional Microscopy at Imperial College London.

## Supplementary Figures

**Fig. S1.**
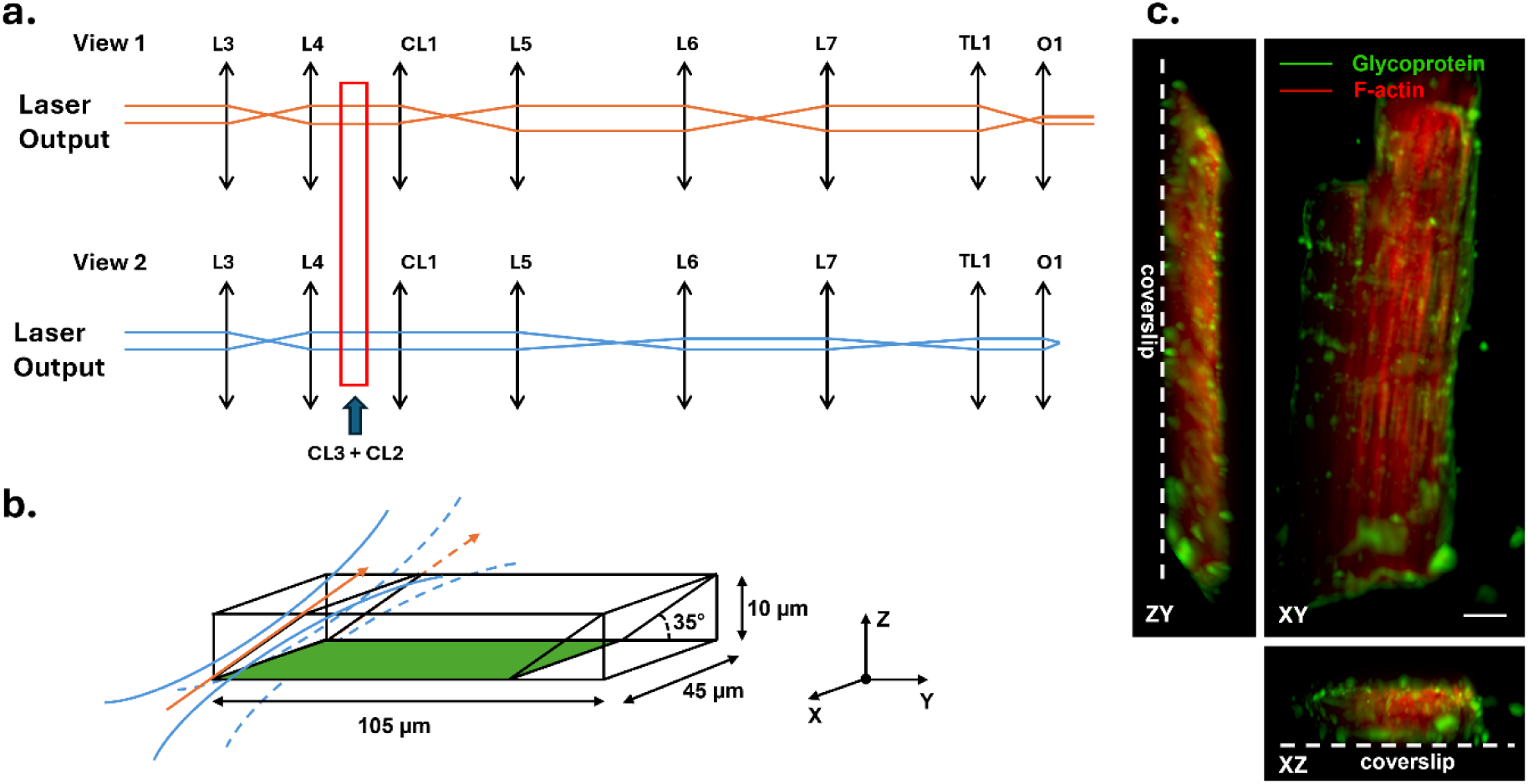
Light sheet shaping and imaging volume of the OPM system. **(a)** Optical ray diagram illustrating the laser beam paths for two orthogonal illumination views (View 1: orange; View 2: blue). After passing through a cylindrical lens (CL1), the beam is focused in one direction while remaining collimated in the orthogonal direction. The subsequent lens elements maintain this anisotropy, forming a thin light sheet at the sample plane. The boxed region (CL2 + CL3) indicates the cylindrical lens pair used to expand the beam laterally. (b) Schematic of the OPM light-sheet scanning volume. The light sheet enters the sample at a 35° angle relative to the coverslip and produces a ∼105 μm × 45 μm × 10 μm imaging volume with high optical sectioning quality. (c) Example volumetric image of a cardiomyocyte stained for F-actin (red) and glycoproteins (green), showing the 3D coverage of the system relative to the coverslip. Scale bar: 10 μm. Cells in (c) were co-stained with WGA-Alexa Fluor™ 488 (10 µg/mL) and CellMask™ Deep Red (1 µM) in high potassium buffer for 30 min at room temperature in the dark. After two washes, samples were imaged immediately.

**Fig. S2.**
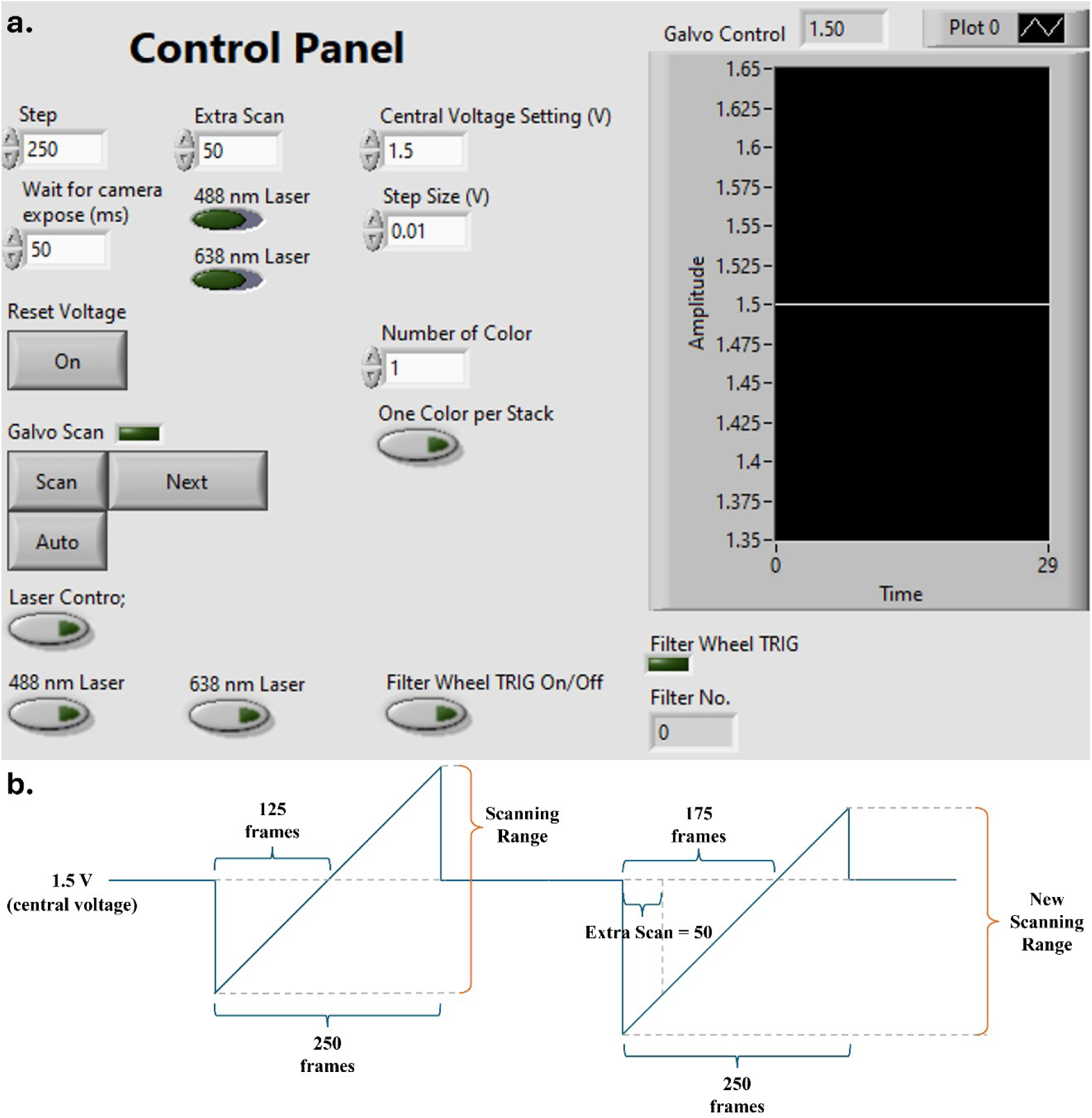
Galvo scanning control interface and scan voltage profile. (a) Custom LabVIEW-based user interface for controlling the galvo mirror during light-sheet scanning. Adjustable parameters include step count, extra scan offset, central voltage, voltage step size, and camera exposure delay. The interface also allows toggling of laser lines (488 nm, 638 nm), filter wheel triggering, and scanning modes. Real-time voltage output is visualised in the “Galvo Control” plot window during acquisition. (b) Schematic of the galvo mirror voltage profile over time. A typical scan consists of 250 steps, with 125 frames collected per direction around a central voltage (e.g., 1.5 V). The “Extra Scan” parameter (e.g., 50) shifts the scanning origin, generating an offset in the scanning range to accommodate asymmetric sample positioning or increase imaging coverage. This flexible scanning profile enables precise alignment and adjustment of the imaged volume.

**Fig. S3.**
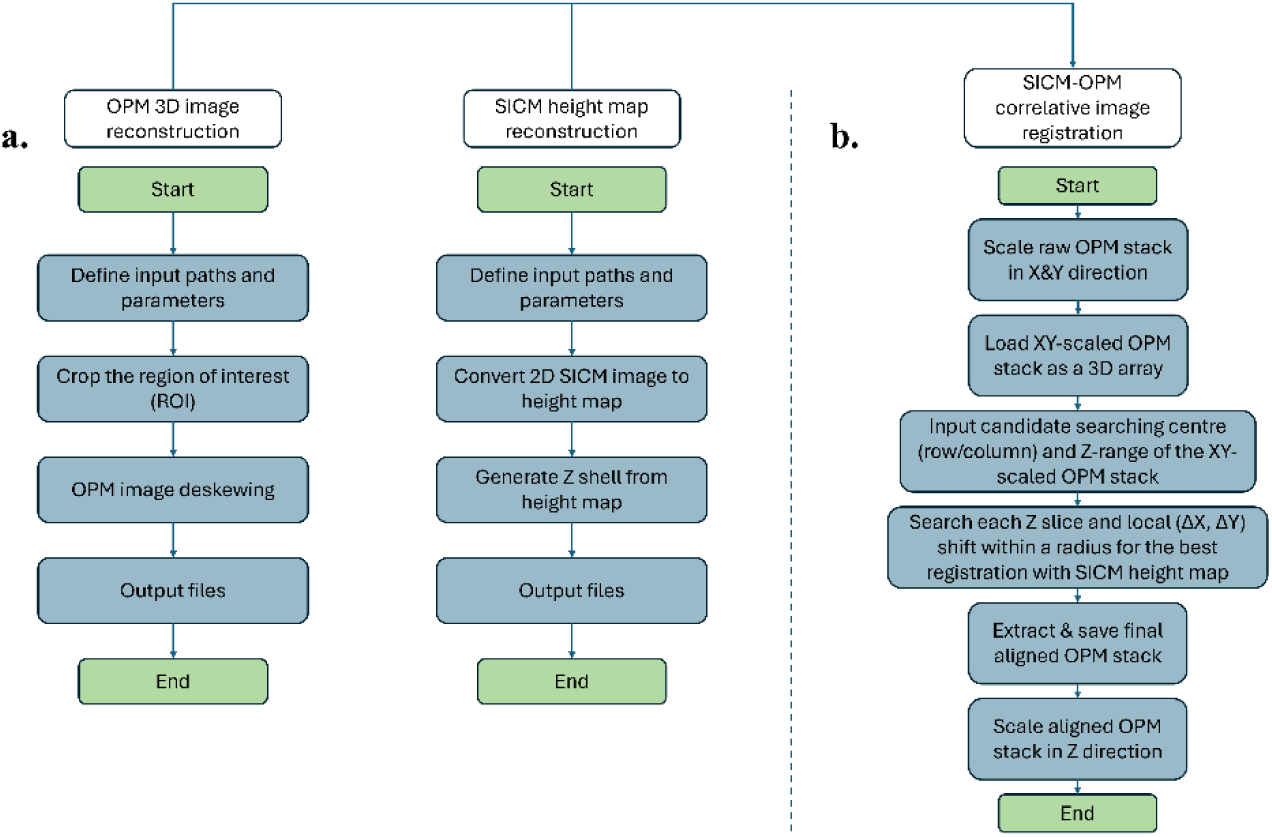
Flowcharts of (a) OPM 3D image reconstruction, SICM height map reconstruction and (b) SICM-OPM correlative image registration.

**Fig. S4.**
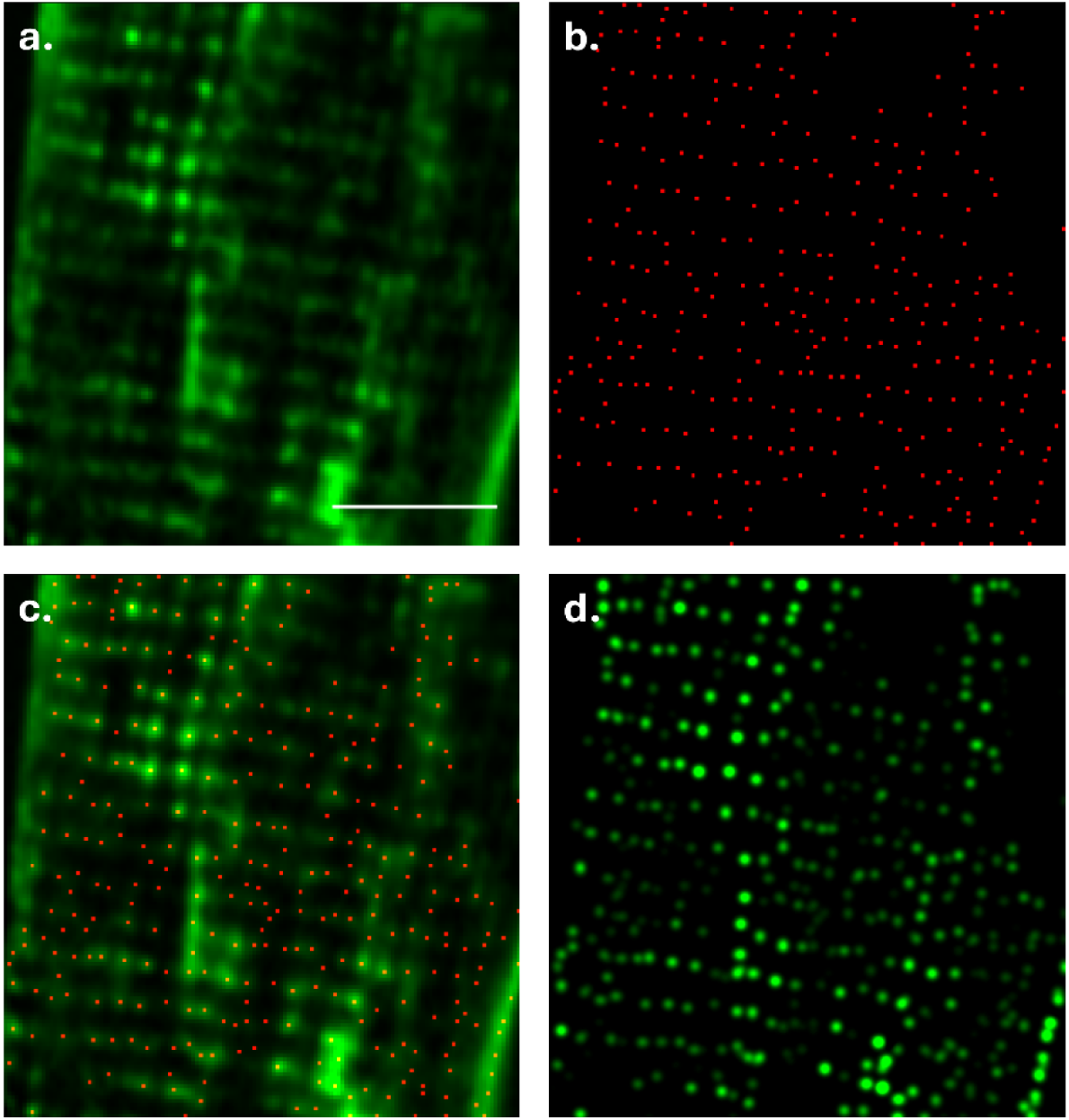
Workflow for generating high-resolution point-based reconstructions from raw fluorescence data. (a) Raw OPM fluorescence image (a single frame) showing periodic subcellular structures. (b) Localisation map of detected features based on Laplacian of Gaussian (LoG) filtering and thresholding; red dots indicate identified peak positions. (c) Overlay of localisations (red) onto the original fluorescence image. (d) High-resolution point-based reconstruction, where each localisation is rendered with a Gaussian kernel onto a high-resolution grid, revealing improved structural clarity. Scale bar: 10 µm.

**Fig. S5.**
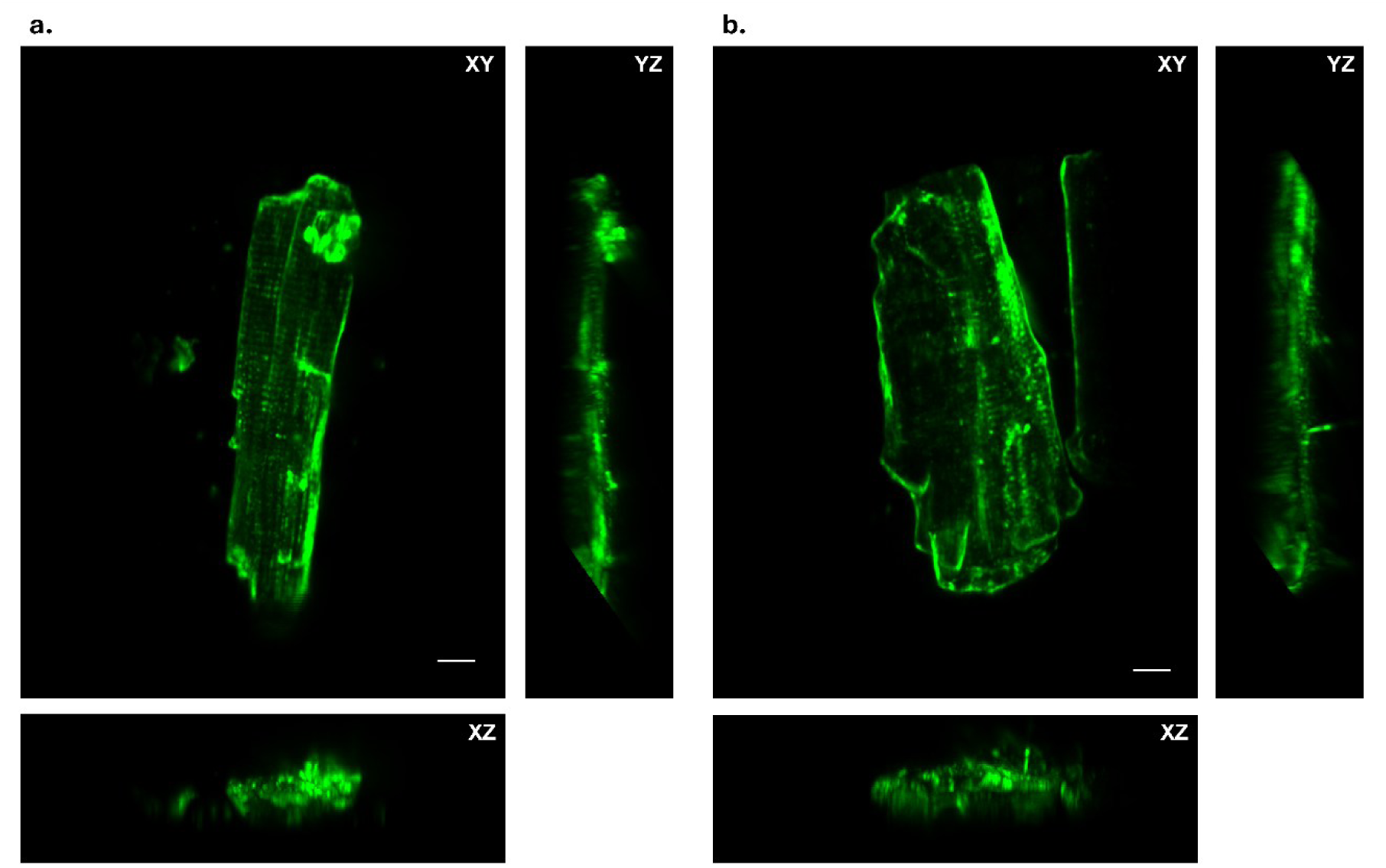
Full-field OPM fluorescence images of control and detubulated cardiomyocytes. (a) Maximum intensity projections (MIPs) of a control adult rat ventricular myocyte (ARVM), displaying well-organised transverse (T)-tubule structures. (b) Corresponding views of an ARVM following imipramine treatment, showing reduced T-tubule signal intensity and disrupted structural continuity, consistent with detubulation. Each panel presents XY, YZ, and XZ MIP views of the same cell. Scale bars: 10 μm.

**Fig. S6.**
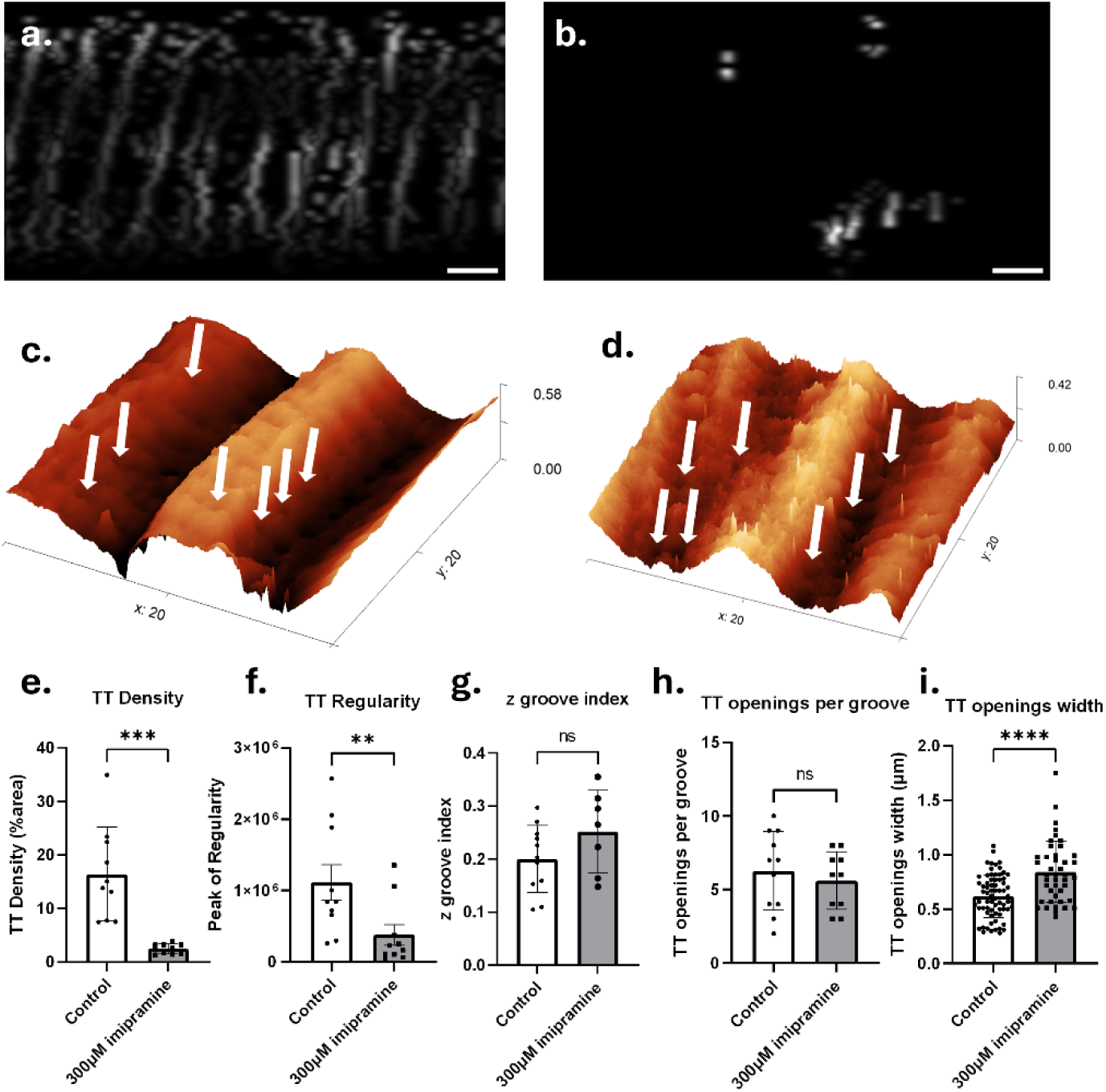
SICM-OPM images and quantitative analysis of T-tubules (TT). (a, b) Representative 2D OPM maximum-intensity projections (2 μm thickness) of Di-8-ANEPPS–labelled adult rat ventricular myocytes (ARVMs) under (a) control and (b) 300 μM imipramine-treated conditions. Images are aligned along the cell’s longitudinal axis to capture a single sarcomere unit. Scale bars: 2 μm. (c, d) Corresponding 3D SICM surface topography images of the same regions in (a, b), highlighting TT openings (white arrows). Axis units are in μm; the vertical height range is shown on the right of each panel. (e, f) Quantification of TT density (control: 16.4% ± 2.8%; imipramine: 2.4% ± 0.3%) and TT regularity (control: 1.11 × 10⁶ ± 2.47 × 10⁵; imipramine: 3.79 × 10⁵ ± 1.43 × 10⁵), corresponding to an 85.1% ± 6.7% reduction in density and 65.9% ± 14.2% decrease in regularity. (g-i) SICM-derived quantification of z-groove index (control: 0.20 ± 0.02; imipramine: 0.25 ± 0.03; with a 25.0% ± 15.6% increase, ns), TT opening number per z-groove (control: 6.27 ± 0.81; imipramine: 5.60 ± 0.62; with a 10.7% ± 14.3% decrease, ns), and TT opening width (control: 0.62 ± 0.02 μm; imipramine: 0.84 ± 0.04 μm; with a 35.5% ± 8.3% significant increase, p<0.0001). Data are presented as mean ± standard error of the mean. Statistical significance: *p < 0.05, **p < 0.01, ***p < 0.001, ****p < 0.0001; ns = not significant. Tests: t-test (e, h, i), Mann– Whitney test (f, g).

**Fig. S7.**
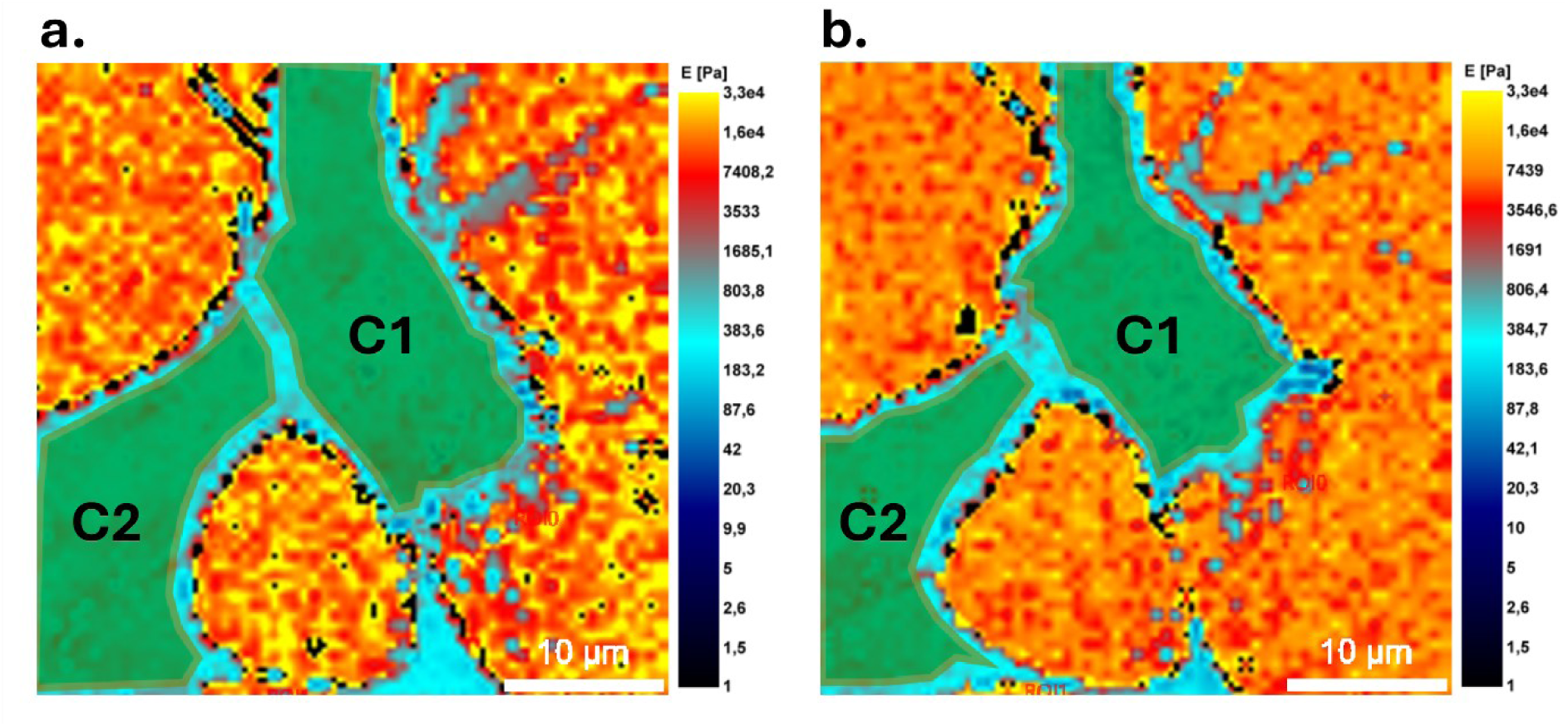
Quantitative comparison of stiffness changes in injected and uninjected SH-SY5Y cells following α-synuclein delivery. (a, b) Young’s modulus (E) maps corresponding to the regions highlighted in Fig. 5(e, f), showing cells C1 (injected) and C2 (uninjected) immediately after injection (a) and 3 hours post-injection (b). At 0 h, the stiffness of C1 and C2 was comparable (C1: 446.0 ± 157.6 Pa; C2: 473.6 ± 150.2 Pa). At 3 h, both cells exhibited softening, but the decrease was more pronounced in the injected cell (C1: 325.6 ± 103.4 Pa; C2: 439.4 ± 142.7 Pa).

